# T cells restrain inflammation driven by TNF-induced cell death

**DOI:** 10.64898/2026.04.21.719867

**Authors:** Michaela Pribikova, Darina Paprckova, Daniela Knizkova, Alzbeta Synackova, Tereza Semberova, Ales Drobek, Veronika Niederlova, Juraj Michalik, Anna Morales Mendez, Ondrej Stepanek, Peter Draber

## Abstract

TNF is a potent proinflammatory cytokine that can induce cell death by activating the kinase RIPK1. The adaptor proteins TANK and AZI2 protect against cell death by recruiting TBK1 to the TNF receptor signaling complex, thereby inhibiting RIPK1. While deficiency of either adaptor alone is well tolerated, combined loss of TANK and AZI2 results in partial embryonic lethality and severe TNF- and RIPK1-driven autoinflammation.

Here, we show that TANK/AZI2-deficient mice exhibit a striking expansion of regulatory T cells (Tregs), most of which display an effector phenotype with high expression of immunosuppressive genes. Although thymic Treg generation is modestly increased, Tregs arise predominantly in the periphery through a largely cell-intrinsic mechanism. The marked accumulation of effector Tregs suggested that the T-cell compartment may limit TNF-driven pathology in this model. Supporting this, T cell ablation in TANK/AZI2-deficient mice markedly exacerbates disease progression and enhances TNF-driven, RIPK1-mediated inflammation. Similarly, T cells protect against acute TNF-induced systemic inflammatory response syndrome by limiting RIPK1-mediated cell death.

Together, our findings identify TANK and AZI2 as negative regulators of Treg formation, demonstrate that T cells restrain TNF-driven inflammation by limiting RIPK1-dependent cell death, and suggest that this protective effect is mediated primarily by Tregs.

## Introduction

Tumor necrosis factor (TNF) is a major proinflammatory cytokine required for immune defense against pathogens, notably *Mycobacterium tuberculosis* (Arias *et al*, 2024). However, aberrant TNF signaling contributes to autoimmune diseases, including rheumatoid arthritis, psoriasis, and inflammatory bowel disease. TNF drives inflammation via the broadly expressed TNF receptor 1 (TNFR1), which can elicit distinct outcomes. In most settings, TNFR1 induces transcriptional responses and the production of proinflammatory mediators. When TNFR1 signaling checkpoints are disrupted, it can trigger apoptotic, pyroptotic, or necroptotic cell death, leading to tissue damage and exacerbated inflammation (van Loo & Bertrand, 2023).

TNF-induced cell death is mediated by receptor-interacting serine/threonine-protein kinase 1 (RIPK1). Following recruitment to the TNFR1 signalosome, RIPK1 can become activated by autophosphorylation, dissociate from the receptor complex, and orchestrate the assembly of a cytosolic cell death-inducing complex II (Micheau & Tschopp, 2003). To suppress complex II formation, RIPK1 activity must be tightly controlled by post-translational modifications, especially ubiquitination and phosphorylation (Clucas & Meier, 2023). One of the kinases that phosphorylates RIPK1 and thereby restrains complex II assembly is TANK-binding kinase 1 (TBK1) (Lafont *et al*, 2018; Xu *et al*, 2018). TBK1 knockout (KO) mice die during embryonic development, which can be rescued by deletion of TNFR1 or inactivation of RIPK1 (Bonnard *et al*, 2000; Eren *et al*, 2024; Xu *et al*., 2018). Human patients with homozygous loss-of-function mutations in TBK1 develop chronic, systemic autoinflammation that can be substantially ameliorated by TNF-blocking antibodies (Taft *et al*, 2021).

Recruitment of TBK1 in the TNFR1 signaling complex is mediated by two adaptors, TANK and AZI2 (also known as NAP1) (Lafont *et al*., 2018). While mice lacking either adaptor are viable and fertile, combined deletion of both proteins results in a severe phenotype (Ujevic *et al*, 2024). *Tank/Azi2* double KO (DKO) mice exhibit partial embryonic lethality, and surviving pups develop multiorgan inflammation, splenomegaly, and die prematurely around six months of age. This phenotype is caused predominantly by TNFR1-mediated cell death, making *Tank/Azi2*^DKO^ mice a valuable genetic model for studying TNF-driven inflammation and the mechanisms that normally prevent it.

Inflammation is tightly controlled by regulatory T cells (Tregs). Tregs require the transcription factor FOXP3 for their development (Fontenot *et al*, 2003; Hori *et al*, 2003; Khattri *et al*, 2003). Loss-of-function mutations in *FOXP3* cause immunodysregulation, polyendocrinopathy, enteropathy, X-linked (IPEX) syndrome (Bennett *et al*, 2001; Borna *et al*, 2024), and Foxp3 deficiency in mice causes severe systemic autoimmunity (Brunkow *et al*, 2001). The core function of Tregs is to suppress autoreactive T cells, since CD3e knockout (KO) mice, which lack both Tregs and conventional T cells, do not develop autoinflammatory disease (Malissen *et al*, 1995). Nevertheless, Tregs regulate diverse immune and non-immune cells to suppress activation of the proinflammatory pathway, promote tissue repair, and limit neuronal activity and pain perception (Dikiy & Rudensky, 2023; Loffredo *et al*, 2024; Mendoza *et al*, 2025). Interestingly, very little is currently known about Tregs’ ability to restrain TNF-driven cell death. Because aberrant activation of cell death is highly proinflammatory, it seems likely that Tregs evolved mechanisms to curb this pathway.

In this work, we found that *Tank/Azi2*^DKO^ mice exhibit markedly increased Treg formation. Surprisingly, ablation of all T cells in *Tank/Azi2/Cd3e* triple KO (TKO) mice substantially worsened autoinflammation. This exacerbated phenotype was rescued by deletion of TNFR1 or inactivation of RIPK1, indicating that T cells, most likely Tregs, protect against TNF-induced cell death. In support of this hypothesis, we show that *Cd3e*^KO^ mice are highly susceptible to injections of otherwise sublethal doses of TNF. Altogether, our data suggest that Tregs can control TNF-driven cell death independently of their canonical role in suppressing autoreactive T cells.

## Results

### *Tank/Azi2*^DKO^ mice display a highly increased proportion of Tregs

The adaptors TANK and AZI2 are remarkably redundant in their ability to prevent autoinflammation. Mice deficient in either adaptor alone do not exhibit an overt phenotype and appear healthy (Ujevic *et al*., 2024). Similarly, mice expressing only one wild-type allele of either adaptor are viable and fertile (Fig. 1A). This allowed us to set up breeding cages to obtain littermate animals that were either heterozygous for both adaptors (control), *Azi2*^+/-^.*Tank*^-/-^(*Tank*^KO^), *Azi2*^-/-^.*Tank*^+/-^ (*Azi2*^KO^), or *Tank/Azi2*^DKO^. Mice lacking both proteins were born in a sub-Mendelian ratio (Fig. 1A) and, at the age of 6-8 weeks, the *Tank/Azi2*^DKO^ mice were smaller and had enlarged spleens compared with their littermates carrying at least one allele of *Tank* or *Azi2* (Fig. 1B).

**Figure 1.**
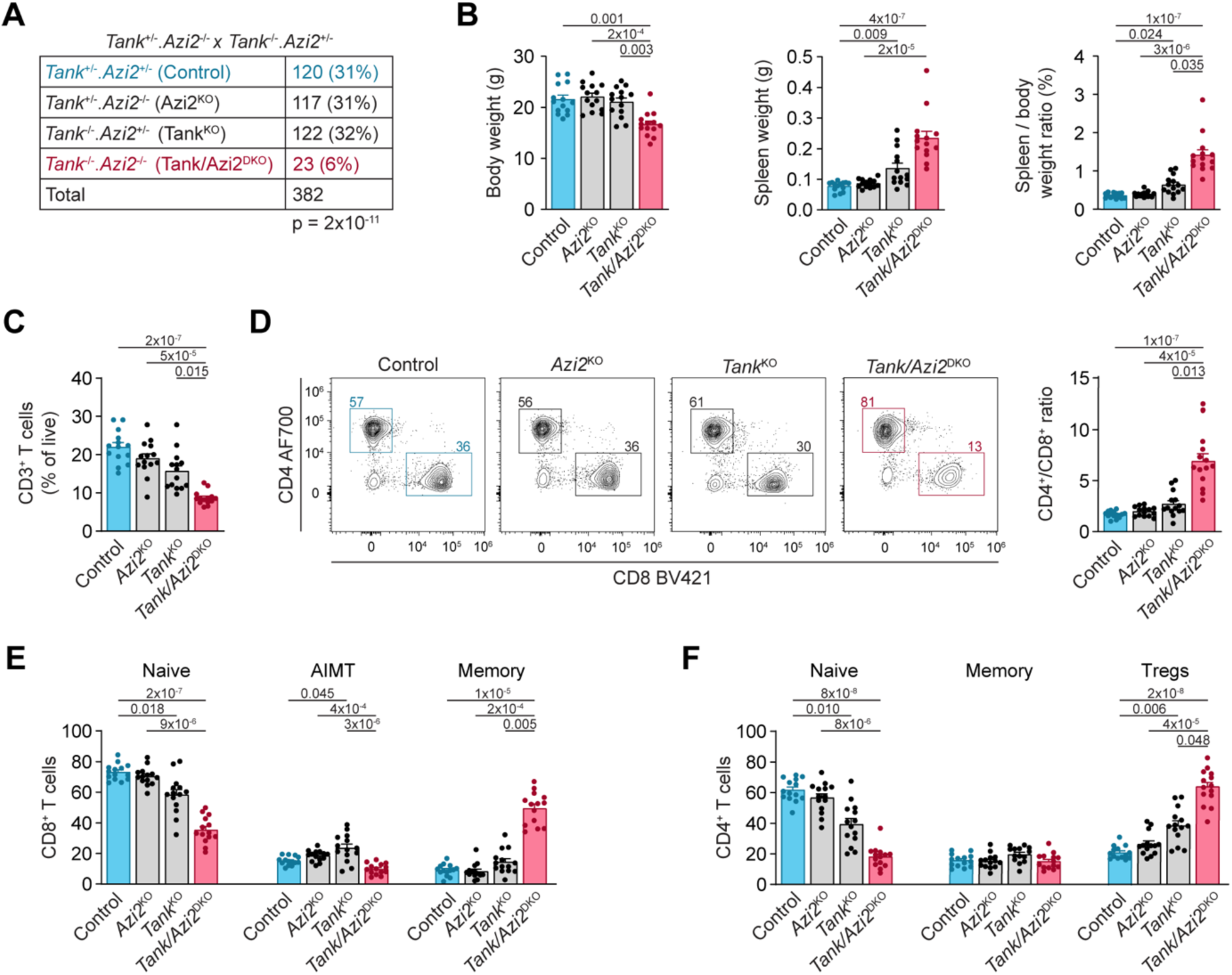
*Tank/Azi2*^DKO^ mice display a highly increased proportion of Tregs. **(A)** Genotypes and numbers of pups born to parents of the indicated genotypes, chi-square test. **(B)** Body weight and spleen weight of 8-12-week-old mice. (**C-F**) Flow cytometry analysis of splenic T cells from 8-12-week-old mice. (C) Proportion of live T cells (gated as CD3^+^). (D) Ratio of CD4^+^ to CD8^+^ T cells. (E) Fraction of CD8^+^ T cells gated as naïve (CD44^−^), memory (CD44^+^, CD49d^+^), and antigen-inexperienced memory T (AIMT) cells (CD44^+^, CD49d^−^). (F) Fraction of CD4^+^ T cells gated as Tregs (FOXP3^+^), naïve (FOXP3^−^, CD44^−^, CD62L^+^), and memory (FOXP3^−^, CD44^+^, CD62L^−^). Data in (B-F) are presented as mean + SEM (n = 14 mice per group), Kruskal–Wallis test with Dunn’s multiple-comparisons post hoc test.

Flow cytometry analysis revealed a decreased proportion of T cells in the spleens of *Tank/Azi2*^DKO^ mice (Fig. 1C, S1A), which is consistent with the previously reported accumulation of isotype-switched B cells (Ujevic *et al*., 2024). The CD4^+^/CD8^+^ T cell ratio was strongly skewed towards CD4^+^ T cells in *Tank/Azi2*^DKO^ animals compared with control mice (Fig. 1D). In accord with ongoing inflammation, the majority of CD8^+^ T cells in Tank/Azi2^DKO^ mice exhibited a memory phenotype (Fig. 1E). In contrast, the analysis of CD4^+^ T cells revealed that the proportion of memory T cells is similar between the strains (Fig. 1F). However, nearly 70% of splenic CD4^+^ T cells in *Tank/Azi2*^DKO^ mice were Tregs, a markedly higher proportion than the approximately 20% observed in control and Azi2^KO^ animals and 40% in *Tank*^KO^ mice (Fig. 1F).

Similarly, the analysis of peripheral lymph nodes showed a decreased proportion of T cells (Fig. S1B), and we observed an increased CD4^+^/CD8^+^ T cell ratio in *Tank/Azi2*^DKO^ mice compared with controls (Fig. S1C). We noted only a small but significant increase in the frequency of conventional memory CD4^+^ and CD8^+^ T cells in lymph nodes of *Tank/Azi2*^DKO^ mice (Fig. S1D-E). Importantly, whereas Tregs accounted for approximately 20% of CD4^+^ T cells in the lymph nodes of control mice, this ratio increased to 40% in *Tank/Azi2*^DKO^ animals (Fig. S1D). Altogether, these results demonstrated that, although *Tank/Azi2*^DKO^ mice develop autoinflammatory disease, they also exhibit a striking increase in Treg frequency in secondary lymphoid organs.

### The majority of Tregs in *Tank/Azi2*^DKO^ mice exhibit an effector phenotype

Next, we aimed to further characterize Tregs in mice lacking the adaptors TANK and AZI2. We performed single-cell RNA sequencing (scRNA-seq) of splenic CD4^+^ T cells from *Tank/Azi2*^DKO^ mice and control littermate *Azi2*^KO^ animals, which do not exhibit an overt phenotype or altered Treg formation. Unsupervised clustering identified 13 clusters, including four Treg clusters characterized by *Foxp3* expression (Fig. 2A-B). Based on the expression of canonical marker genes, we annotated Treg clusters as naive, transient, effector, and RORγt^+^ Tregs. Conventional CD4^+^ T cell clusters were annotated as naive, T helper type 17 (Th17), T follicular helper (Tfh), interferon-stimulated gene (ISG)-high, early activated, cytotoxic-leaning, and cytotoxic. We also detected a cluster of proliferating cells (Fig. 2C).

**Figure 2.**
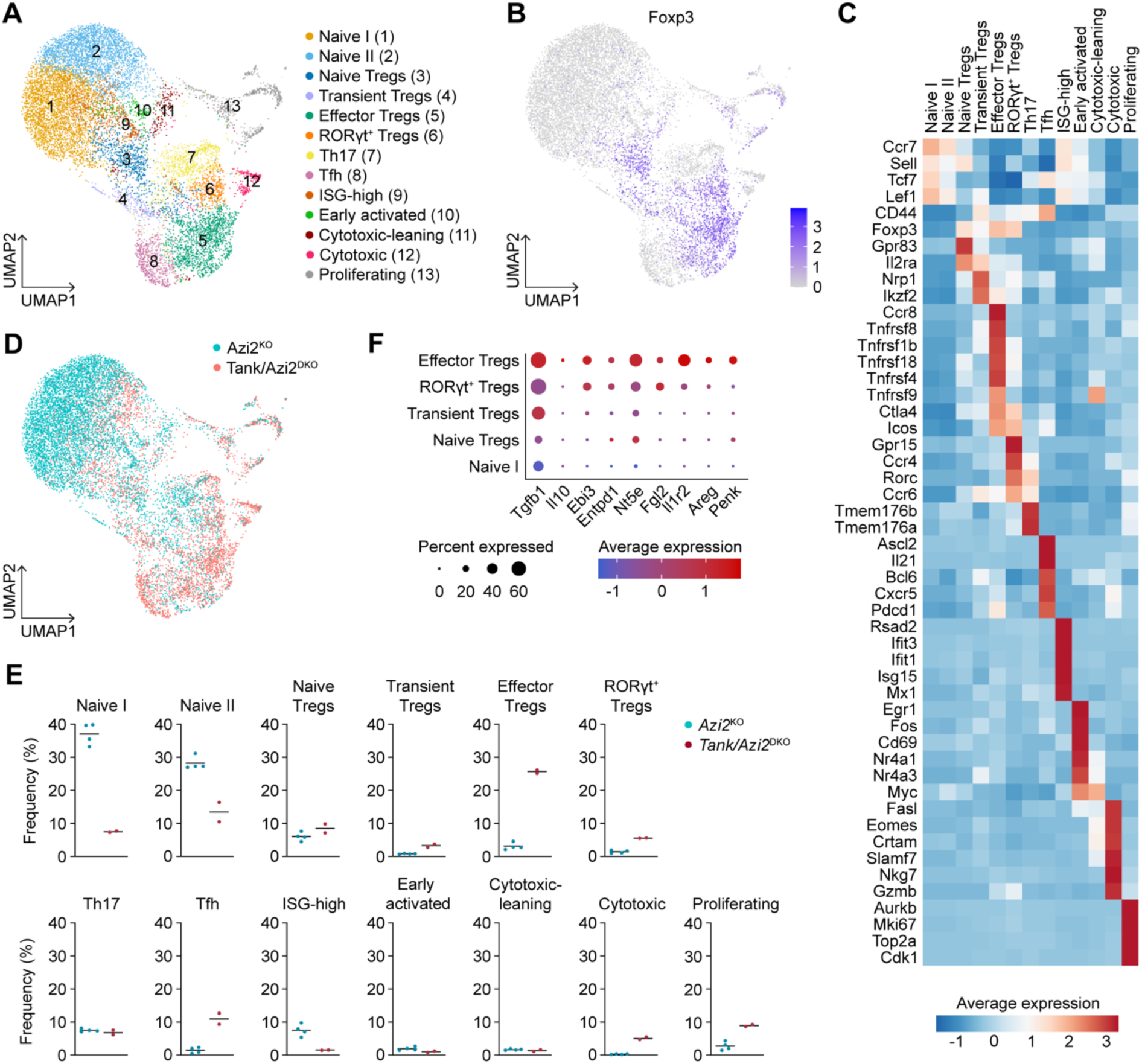
The majority of Tregs in *Tank/Azi2*^DKO^ mice exhibit an effector phenotype. scRNA-seq analysis of splenic CD4^+^ T cells isolated from *Azi2*^KO^ (n = 4) and *Tank/Azi2*^DKO^ mice (n = 2). **(A)** UMAP plot of individual cells with unsupervised clustering and manual cluster annotation. **(B)** UMAP plot showing expression of the Treg marker *Foxp3*. **(C)** Heatmap of marker genes defining the individual clusters. **(D)** UMAP plot showing the distribution of cells of the indicated genotype **(E)** Proportion of cells in the different clusters plotted as a percentage of all cells from each genotype. **(F)** Dot plot showing expression of the indicated immunosuppressive genes in Treg clusters.

The majority of CD4^+^ T cells in control *Azi2*^KO^ mice displayed a naive transcriptional signature. In contrast, *Tank/Azi2*^DKO^ mice showed a substantially reduced proportion of naive cells (Fig. 2D-E). This shift was accompanied by an increased proportion of Tfh cells, consistent with expansion of memory B cells in these mice (Ujevic *et al*., 2024), and an increased proportion of cytotoxic CD4^+^ T cells, which are often enriched during chronic viral infections and also in several autoimmune diseases (Oh & Fong, 2021). Cells from *Tank/Azi2*^DKO^ mice were enriched in all Treg clusters, most notably in effector Tregs, which represented approximately 5% of all splenic CD4^+^ T cells in *Azi2*^KO^ mice but nearly 25% in *Tank/Azi2*^DKO^ mice (Fig. 2D-E). Subsets enriched in *Tank/Azi2*^DKO^ mice also showed higher clonal abundance by V(D)J sequencing (Fig. S2A), in line with greater CD4^+^ T cell clonal expansion observed in the *Tank/Azi2*^DKO^ animals (Fig. S2B).

The effector Treg cluster exhibited high expression of *Ccr8*, a hallmark of a highly immunosuppressive population found in inflamed tissues and cancer (Barsheshet *et al*, 2017; Kidani *et al*, 2022). Consistent with this, we detected increased expression of established immunoregulatory genes (Fig. 2F). These included anti-inflammatory cytokines *Tgfb1*, *Il10*, or IL-35 subunit *Ebi3*, the ectoenzymes *Entpd1* (CD39) and *Nt5e* (CD73), which cooperate to convert extracellular ATP into anti-inflammatory adenosine, immunosuppressive factors *Fgl2* and the IL-1 decoy receptor *Il1r2*, the tissue repair factor *Areg*, and the neuromodulator *Penk* (Arnouk *et al*, 2025; Dikiy & Rudensky, 2023; Mendoza *et al*., 2025; Shalev *et al*, 2008).

To corroborate our findings, we took advantage of the fact that effector Tregs are characterized by high expression of several TNF receptor superfamily members (Vasanthakumar *et al*, 2017). Using flow cytometry, we demonstrated that splenic Tregs from *Tank/Azi2*^DKO^ mice exhibited increased expression of effector Treg markers, including GITR (TNFRSF18), OX-40 (TNFRSF4), 4-1BB (TNFRSF9), as well as TNFR2 (TNFRSF1B) (Fig. S2C-D). Altogether, these data indicate that *Tank/Azi2*^DKO^ mice exhibit a striking expansion of effector Treg cells.

### *Tank/Azi2*^DKO^ CD4^+^ T cells have an intrinsic propensity to form Tregs at the periphery

To determine whether the increased proportion of Tregs in *Tank/Azi2*^DKO^ mice is due to skewed thymic development, we isolated thymi from 5-week-old control *Azi2*^KO^ mice and compared them with those from *Tank/Azi2*^DKO^ littermates. The generation of CD4^+^/CD8^+^ double-positive thymocytes was not altered. Similarly, the emergence of CD4^+^ and CD8^+^ single-positive thymocytes did not differ significantly between the two strains (Fig. 3A), and their ratio was not significantly changed (Fig. 3B). However, Tregs constituted, on average, around 20% of CD4^+^ single-positive thymocytes in *Tank/Azi2*^DKO^ mice, a proportion nearly threefold higher than in control *Azi2*^KO^ littermates (Fig. 3C).

**Figure 3.**
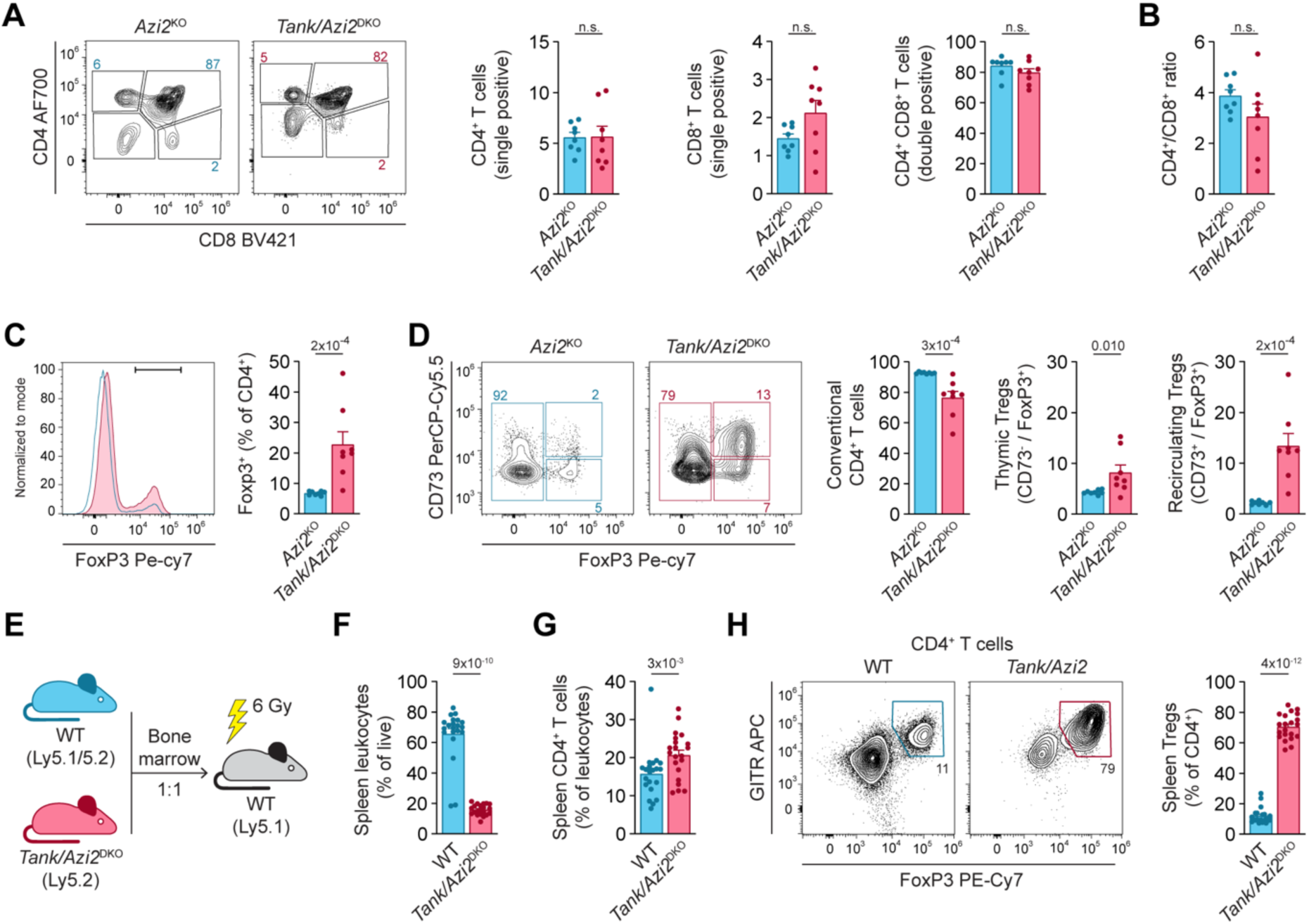
*Tank/Azi2*^DKO^ CD4^+^ T cells have an intrinsic propensity to form Tregs in the periphery. (**A-D**) Flow cytometry analysis of thymocytes from 5-week-old *Azi2*^KO^ and *Azi2/Tank*^DKO^ (n = 8 mice per group) mice. (A) T cells (CD3^+^) were gated as CD4/CD8 double-positive, and CD4 or CD8 single-positive T cells. (B) Ratio between CD4 and CD8 single-positive thymocytes. (C) Proportion of Tregs among CD4 single-positive T cells. (D) CD4 single-positive T cells were gated as conventional CD4^+^ T cells (FOXP3^−^), thymic Tregs (FOXP3^+^, CD73^−^), and circulating Tregs (FOXP3^+^, CD73^+^). (**E-H**) Flow cytometry analysis of splenic T cells isolated from irradiated recipient mice reconstituted with mixed bone marrow chimeras generated from WT and *Azi2/Tank*^DKO^ donors (n = 21 recipient mice in 3 separate experiments). (E) Experimental scheme allowing the donor cells to be distinguished by the congenic markers Ly5.1 and Ly5.2. (F) Percentage of splenic leukocytes derived from the indicated bone marrow donor cells. (G) Percentage of CD4⁺ T cells among splenic leukocytes. (H) Frequency of Tregs (GITR^high^, FOXP3⁺) within the CD4⁺ T cell population. Data are shown as mean + SEM. Two-tailed Mann-Whitney test. n.s., not significant.

About half of the thymic Tregs in wild-type mice are mature, recirculating cells that can be distinguished by high CD73 expression (Owen *et al*, 2019; Voboril *et al*, 2020) and we observed a similar ratio in control *Azi2*^KO^ mice (Fig. 3D). Notably, the fraction of recirculating Tregs was highly increased in the thymi of *Tank/Azi2*^DKO^ animals, while we detected only modest increase in de novo thymic Treg generation (Fig. 3D). These data suggest that the massive expansion of Tregs in *Tank/Azi2*^DKO^ mice occurs predominantly in peripheral lymphoid organs, and subsequently these cells recirculate back to the thymus.

To determine whether TANK and AZI2 restrain Treg formation in a cell-intrinsic manner, we performed competitive bone marrow transfer. Bone marrow cells from WT and *Tank/Azi2*^DKO^ mice were mixed at a 1:1 ratio and used to reconstitute irradiated WT recipients. Cells from each donor strain were distinguished by the congenic markers Ly5.1 and Ly5.2 (Fig. 3E). Flow cytometry revealed that WT bone marrow cells engrafted more efficiently than *Tank/Azi2*^DKO^ cells. However, the proportion of splenic CD4^+^ T cells among leukocytes was similar between the two donors (Fig. 3F-G, S3A-B). Importantly, nearly 80% of *Tank/Azi2*^DKO^ splenic CD4^+^ T cells were Tregs, while this ratio was approximately 10% among CD4^+^ T cells derived from WT bone marrow (Fig. 3H). Similar results were observed in lymph nodes, where nearly 60% of *Tank/Azi2*^DKO^ CD4^+^ T cells were Tregs (Fig. S3A-C). Altogether, these data indicate that *Tank/Azi2*^DKO^ CD4^+^ T cells preferentially generate large numbers of Tregs in a cell-intrinsic manner.

### *Tank/Azi2*^DKO^ mice exhibit increased Treg formation in the absence of TNFR1-driven inflammation

An increased emergence of effector Tregs has been reported in inflamed tissues, including arthritic joints, where TNF-mediated tissue destruction plays an important role (Mijnheer *et al*, 2021; Schnell *et al*, 2025). Because *Tank/Azi2*^DKO^ mice develop TNFR1-driven autoinflammation that is largely mediated by RIPK1-dependent cell death (Ujevic *et al*., 2024), we asked whether the ongoing inflammatory environment contributes to the enhanced accumulation of Treg cells.

We analyzed splenic T cells isolated from *Tank/Azi2/Tnfr1* triple KO (TKO) mice, which do not develop autoinflammatory disease, and compared them with *Azi2/Tnfr1*^DKO^ littermates. Although we observed a decreased proportion of splenic T cells in Tank/*Azi2/Tnfr1*^TKO^ animals (Fig. 4A), the ratio between CD4^+^ and CD8^+^ T cells was only mildly increased (Fig. 4B). Notably, nearly 40% of splenic CD4^+^ T cells in Tank/*Azi2/Tnfr1*^TKO^ mice were Tregs, compared with approximately 20% in control animals (Fig. 4C). In contrast, we did not detect substantial increase in memory CD4^+^ T cells (Fig. 4C). We obtained similar results in the lymph nodes (Fig. S4A-C). Because murine T cells express little or no TNFR1, especially when compared to myeloid cells (Fig. S4D), it seems unlikely that loss of TNFR1 directly alters T cell development. These data indicate that removing TNFR1 in *Tank/Azi2*^DKO^ mice abolishes the inflammatory environment and prevents the accumulation of memory T cells, while still permitting robust Treg enrichment.

**Figure 4.**
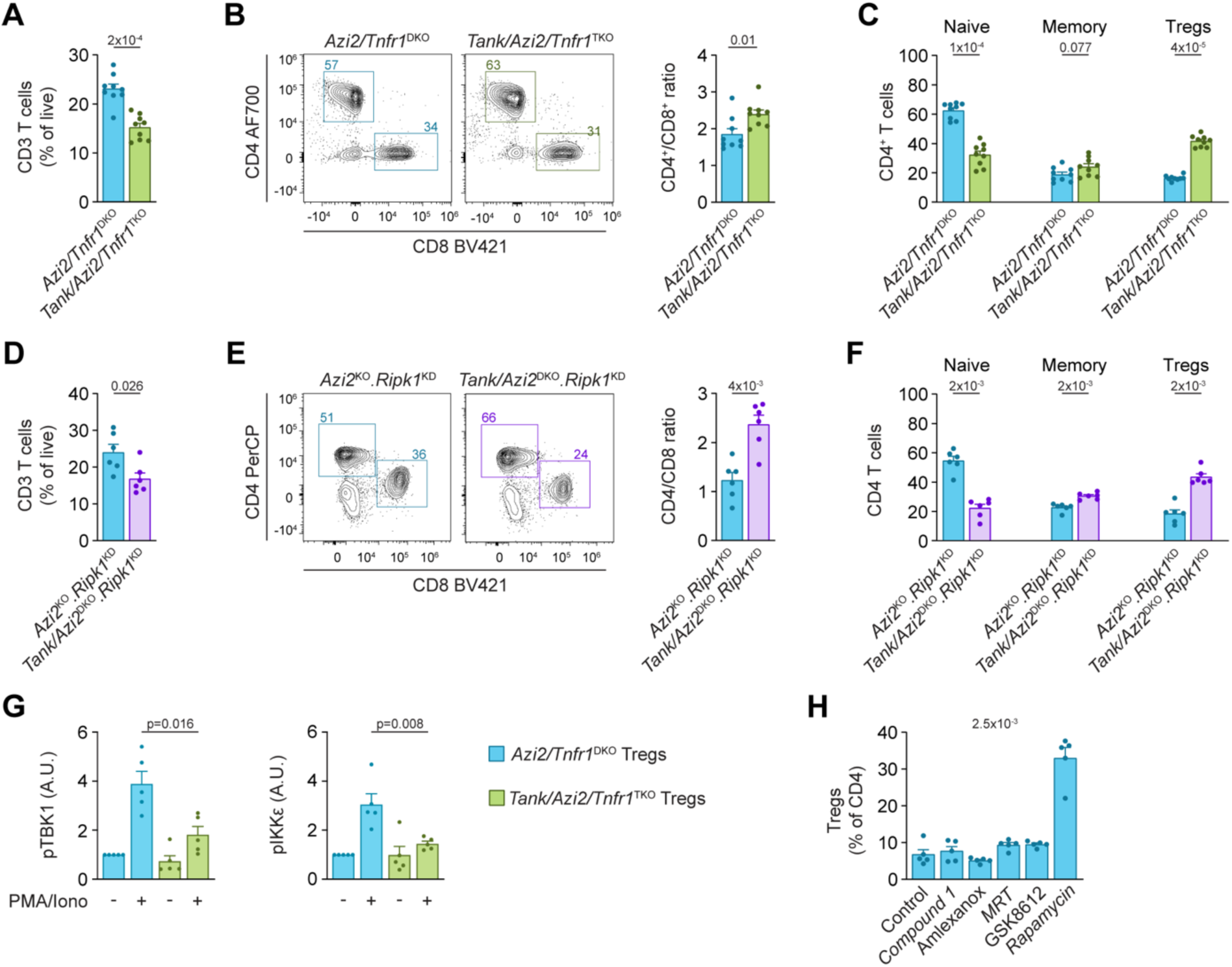
Enhanced Tregs formation in *Tank/Azi2*^DKO^ mice is partially driven by TNFR1-mediated inflammation. (**A-F**) Flow cytometry analysis of splenic T cells from 8-12-week-old *Azi2/Tnfr1*^DKO^ and *Tank/Azi2/Tnfr1*^TKO^ mice (n = 9 mice per group) (A-C) or *Azi2*^KO^.*Ripk1*^KD^ and *Tank/Azi2*^DKO^.*Ripk1*^KD^ mice (n = 6 mice per group) (D-F). (A, D) Proportion of live T cells (gated as CD3^+^). (B, E) Ratio of CD4^+^ to CD8^+^ T cells. (C, F) Fraction of CD4^+^ T cells gated as Tregs (FOXP3^+^), naïve (FOXP3^−^, CD44^−^, CD62L^+^) and memory cells (FOXP3^−^, CD44^+^, CD62L^−^). **(G)** Splenocytes from *Azi2/Tnfr1^DKO^*and *Tank/Azi2/Tnfr1^TKO^* mice were stimulated with PMA/ionomycin for 15 min. Phosphorylation of TBK1 and IKKε in Tregs was analyzed by flow cytometry in 5 independent experiments. **(H)** Splenic T cells were stimulated with anti-CD3/CD28 beads in the presence of IL-2 and the indicated inhibitors for 5 days. The proportion of Tregs among CD4^+^ T cells was analyzed by flow cytometry in 5 independent experiments. Data are presented as mean + SEM. Two-tailed Mann-Whitney test (A-G), or Kruskal–Wallis test (H).

To further support these findings, we analyzed the immune system in *Tank/Azi2*^DKO^ mice carrying the RIPK1 kinase-dead (KD) D138N mutation, which largely prevents TNF-induced cell death and substantially ameliorates disease in *Tank/Azi2*^DKO^ animals (Polykratis *et al*, 2014; Ujevic *et al*., 2024). We again observed an increased proportion of Tregs in both spleen (Fig. 4D-F) and lymph nodes (Fig. S4E-G). Altogether, our results suggest that CD4^+^ T cells in *Tank/Azi2*^DKO^ mice are intrinsically skewed towards Treg differentiation.

### Enhanced formation of Tregs in *Tank/Azi2*^DKO^ mice is not caused by impaired TBK1 and IKKε **activation**

The adaptor proteins TANK and AZI2 promote the activation of TBK1 and IKKε in response to various stimuli, including TNF or IL-17A (Draberova *et al*, 2020; Lafont *et al*., 2018; Ujevic *et al*., 2024). We therefore hypothesized that enhanced Treg formation in *Tank/Azi2*^DKO^ mice might result from defective activation of TBK1 and/or IKKε in T cells. To avoid confounding effects of autoinflammation on immune composition and basal signaling, we isolated splenocytes from *Tank/Azi2/Tnfr1*^TKO^ mice, which do not develop disease, and from control *Azi2/Tnfr1*^DKO^ littermates. Cells were stimulated with phorbol 12-myristate 13-acetate (PMA) and ionomycin, a treatment that mimics strong downstream TCR pathway activation (Lee *et al*, 2023). This treatment induced robust phosphorylation of TBK1 and IKKε in control Tregs, which was substantially reduced in cells lacking both TANK and AZI2 (Fig. 4G), indicating that these adaptors contribute to TBK1 and IKKε activation in this context.

We next tested whether reduced TBK1 and IKKε activity could account for the increased propensity of naive CD4^+^ T cells to differentiate into Tregs. WT splenocytes were stimulated with anti-CD3ε/anti-CD28 antibodies and IL-2 in the presence of various inhibitors. As shown previously, the mTOR inhibitor rapamycin strongly promoted Treg emergence (Battaglia *et al*, 2005; Sauer *et al*, 2008). In contrast, none of the TBK1/IKKε inhibitors tested, including Compound 1, amlexanox, MRT67307, or GSK8612 (Clark *et al*, 2011; Jenkins *et al*, 2018; Reilly *et al*, 2013; Thomson *et al*, 2019), increased Treg formation (Fig. 4H).

These results suggest that enhanced Treg formation caused by loss of TANK and AZI2 is unlikely to be mediated solely by reduced TBK1 and IKKε activity. Consistent with this conclusion, a recent study reported that although *TBK1/IKKε* ^DKO^ mice are embryonically lethal, *Tbk1/Ikke*^DKO^.*Ripk1*^KD^ animals survive to adulthood. However, these mice do not show an increased proportion of splenic Tregs, in contrast to *Tank/Azi2*^DKO^.*Ripk1*^KD^ animals (Eren *et al*., 2024).

### T cell ablation exacerbates TNF-driven autoinflammatory disease in *Tank/Azi2*^DKO^ mice

The increased propensity of T cells to form Tregs in *Tank/Azi2*^DKO^ mice led us to question their role in driving autoinflammation in this model. We removed all T cells by crossing *Tank/Azi2*^DKO^ mice with CD3ε-deficient animals, which lack a crucial subunit of the TCR complex required for thymic T cell development (DeJarnette *et al*, 1998).

*Tank/Azi2/Cd3e*^TKO^ mice were born at a similarly sub-Mendelian ratio as *Tank/Azi2*^DKO^ mice (Fig. S5A). Both strains displayed delayed hair growth at 3 weeks of age (Fig. 5A). However, the *Tank/Azi2/Cd3e*^TKO^ animals were significantly smaller (Fig. 5B) and died around 4 weeks of age, while *Tank/Azi2*^DKO^ mice had a median survival of approximately 25 weeks (Fig 5C). Histological examination revealed markedly increased swelling of glomeruli in the kidneys of *Tank/Azi2/Cd3e*^TKO^ mice, indicating exacerbated glomerular injury (Fig. 5D). These results indicated that T cells alleviate the autoinflammatory disease in *Tank/Azi2*^DKO^ mice.

**Figure 5.**
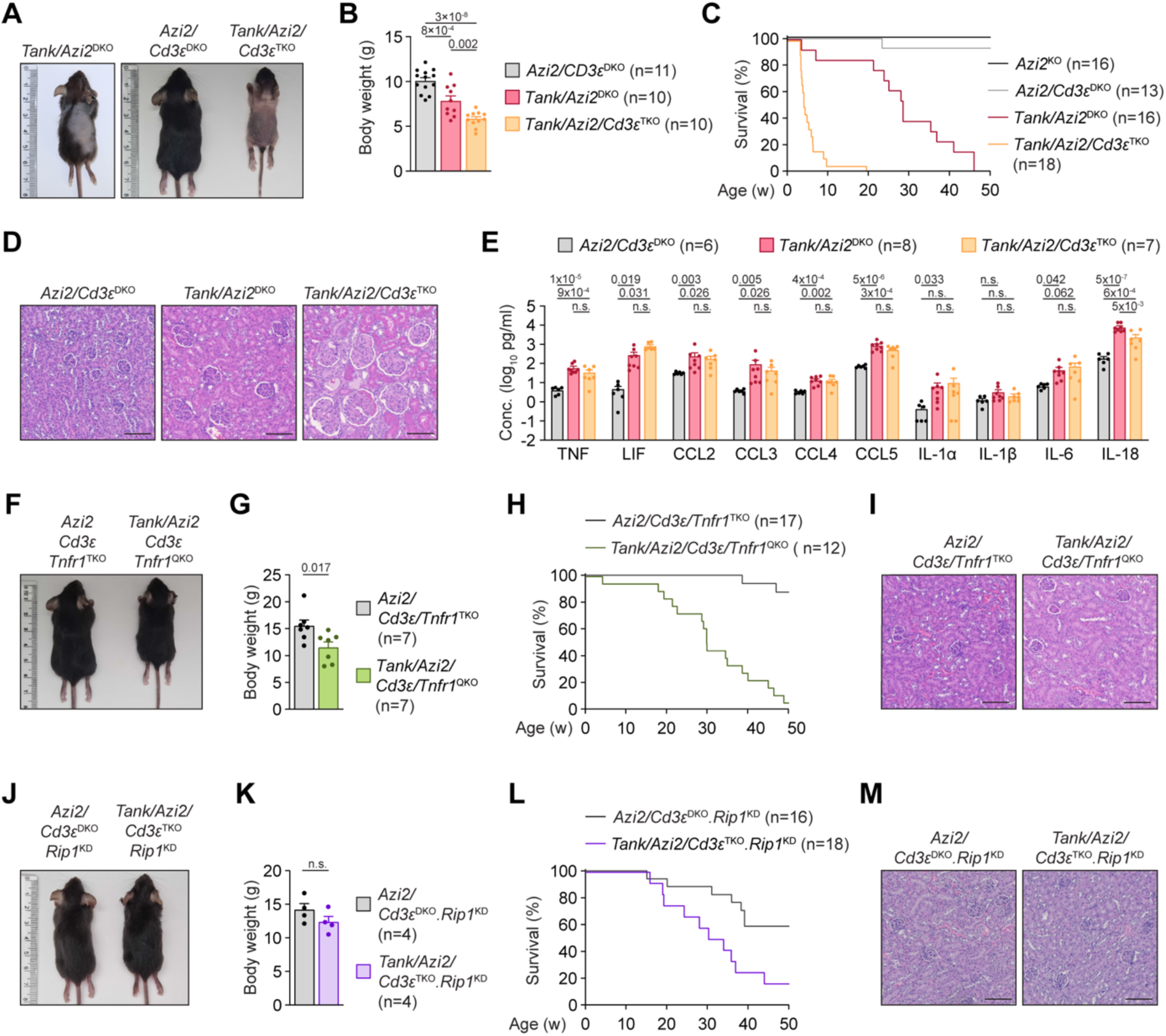
T cell ablation exacerbates TNF-driven autoinflammatory disease in *Tank/Azi2*^DKO^ mice. (**A-E**) Analysis of *Tank/Azi2*^DKO^*, Tank/Azi2/CD3e*^TKO^ and control *Azi2/CD3e*^DKO^ mice. (A) Representative photographs and (B) body weight of 22-26-day-old mice. (C) Survival curves. (D) H&E-stained kidney sections from 22-26-day-old mice, representative images from 5 mice. (E) Plasma cytokine concentrations in 22-26-day-old mice. (**F-K**) Rescue of *Tank/Azi2/Cd3e*^TKO^ mice by *Tnfr1*^KO^ or *Ripk1*^KD^ alleles. (F, J) Representative photographs and (G, K) body weight of 25-33-day-old mice. (H, L) Survival curves. (I, M). H&E-stained kidney sections from 25-33-day-old mice, representative images from 4 mice. Graphs (B, E, G, K) are shown as mean + SEM. one-way ANOVA with Tukey’s post-tests (B, E), Two-tailed Mann-Whitney test (G, K). n.s., not significant. The scale bar is 100 µm.

The most likely explanation for this unexpected phenotype is that the absence of T cells also results in the loss of Tregs. Tregs have previously been shown to suppress the production of proinflammatory cytokines, including TNF, in various immune cell types (Ou *et al*, 2023). We therefore compared the serum cytokine profile of *Tank/Azi2/Cd3e*^TKO^ mice with that of *Tank/Azi2*^DKO^ and control animals at 3 weeks of age. Although the concentrations of several highly proinflammatory cytokines, including TNF, were increased in *Tank/Azi2*^DKO^ mice, their levels were not further increased in *Tank/Azi2/Cd3e*^TKO^ animals (Fig. 5E and S5B), indicating that the accelerated disease and elevated kidney damage in these mice are not simply caused by enhanced systemic production of proinflammatory cytokines.

Next, we aimed to determine whether the early-onset disease in *Tank/Azi2/Cd3e*^TKO^ mice is caused by enhanced TNF-driven inflammation. Indeed, *Tank/Azi2/Cd3e/Tnfr1* quadruple KO (QKO) mice were born at a normal ratio (Fig. S5C) and, at 3 weeks of age, had normal hair growth, although they were slightly smaller (Fig. 5F-G). Most importantly, their median survival was prolonged to approximately 25 weeks (Fig. 5H), and glomerular swelling in the kidneys was rescued (Fig. 5I). We observed a similar rescue upon crossing *Tank/Azi2/Cd3e*^TKO^ to *Ripk1*^KD^ (Fig. 5J-M and S5D). Because RIPK1 kinase activity is required for TNF-induced cell death, these data indicate that aberrant cell death is a major driver of inflammation in this model. Altogether, these results suggest that T cells, most likely Tregs, suppress TNF-induced and RIPK1-dependent cell death in *Tank/Azi2*^DKO^ mice.

### T cells protect against TNF-induced cell death

To evaluate whether T cells can prevent TNF-induced cell death, we used a model of acute TNF-induced inflammation. Injection of mice with a high dose of TNF induces an acute systemic inflammatory response syndrome (SIRS), which is characterized by hypothermia and organ failure. SIRS depends on TNFR1 and requires RIPK1 kinase activity (Duprez *et al*, 2011; Polykratis *et al*., 2014). Since our data indicated that T cells protect from TNF-mediated inflammation in *Tank/Azi2*^DKO^ mice, we aimed to evaluate whether they restrain RIPK1-dependent cell death in the SIRS model.

Injection of *Cd3e*^KO^ mice with a sublethal dose of TNF caused a rapid drop in body temperature, and most animals succumbed to TNF administration within 12 hours. Importantly, hypothermia and lethality were prevented by administration of RIPK1 inhibitor Necrostatin-1s (Nec-1s) prior to TNF injection (Fig. 6A-B). Similarly, *Cd3e*^KO^.*Ripk1*^KD^ mice harboring kinase-dead RIPK1 were protected from TNF challenge, while littermate *Cd3e*^KO^ mice were highly sensitive (Fig. 6C-D). These data showed that T cells protect against TNF-induced SIRS by suppressing RIPK1 kinase activation. In accord, it has previously been shown that *Rag*^KO^ mice lacking both T and B cells exhibit exacerbated hypothermia upon TNF administration compared with control mice (Muendlein *et al*, 2022), a finding we also confirmed (Fig. S6A-B). Taken together with our previous results, these findings support the conclusion that T cells, probably Tregs, can inhibit TNF-induced, RIPK1-dependent cell death, which likely explains their role in slowing TNF-driven disease progression in *Tank/Azi2*^DKO^ mice.

**Figure 6.**
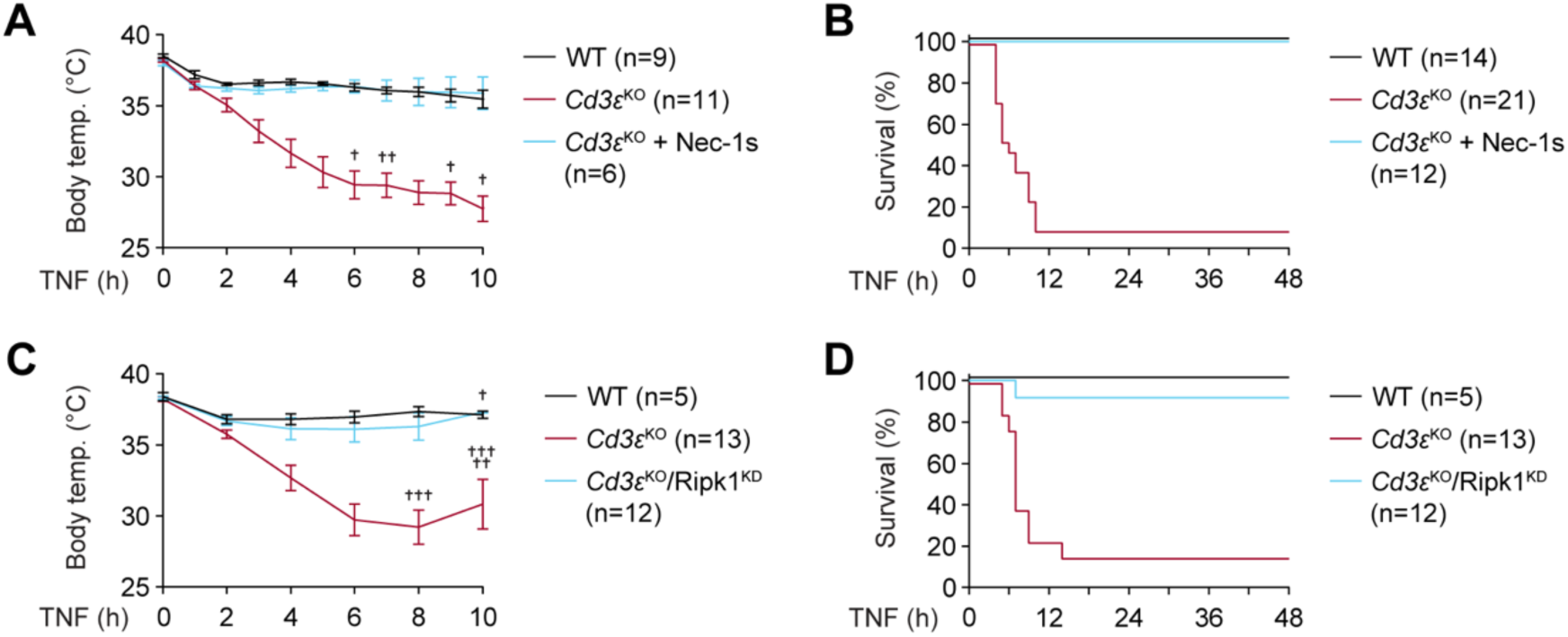
T cells protect against TNF-induced cell death. (**A-B**) WT and *Cd3e*^KO^ mice were injected intraperitoneally with Nec-1s (10 µg/g body weight) in DMSO or with vehicle alone. After 30 minutes, mice were injected intravenously with TNF (1 µg/g body weight). (A) Body temperature was measured every hour for up to 10 h after TNF administration. **(B)** Proportion of mice reaching humane endpoint. (**C-D**) WT or *Cd3e*^KO^, and *Cd3e/Ripk1*^DKO^ littermates were injected with TNF (1 µg/g body weight). (C) Body temperature was measured every 2 h for up to 10 h. (D) Proportion of mice reaching humane endpoint. The mice that reached humane endpoint are indicated by a cross.

## Discussion

This study identified an unexpected feature of *Tank/Azi2*^DKO^ mice. While mice expressing a single allele of either adaptor remain healthy and fertile, combined deficiency of both proteins results in TNFR1-driven multiorgan autoinflammation, accompanied by partial embryonic lethality, premature mortality, and increased emergence of isotype-switched B cells (Ujevic *et al*., 2024). At the same time, we discovered that nearly 60% of splenic CD4^+^ T cells in *Tank/Azi2*^DKO^ mice are Tregs. This surprising finding prompted us to characterize Tregs in this model and to investigate the role of T cells in regulating TNFR1-driven pathology.

Treg development in the thymus was only modestly increased in *Tank/Azi2*^DKO^ mice, suggesting that the major Treg expansion occurs in the periphery. Consistent with this, most splenic Tregs in these mice were clonally expanded and displayed an effector phenotype with elevated expression of CCR8, a marker of highly suppressive activated Tregs (Wen *et al*, 2025). Competitive bone marrow transplantation further demonstrated that the increased propensity to generate Tregs is largely cell-intrinsic. However, inflammation appears to further promote their accumulation, as nearly 70% of splenic CD4^+^ T cells were Tregs in *Tank/Azi2*^DKO^ mice, whereas this proportion decreased to 40% in *Tank/Azi2/Tnfr1*^TKO^ mice and *Tank/Azi2*^DKO^.*Ripk1*^KD^, which do not develop autoinflammation.

The molecular mechanisms underlying enhanced Treg formation in *Tank/Azi2*^DKO^ mice remain unclear. One possible explanation is increased activation of the NF-κB signaling pathway. Indeed, *Tank*^KO^ and *Tank/Azi2*^DKO^ cells exhibit enhanced NF-κB activation downstream of multiple receptors, including Toll-like receptors, the IL-1 receptor, and TNFR1 (Kawagoe *et al*, 2009; Ujevic *et al*., 2024). In T cells, enforced NF-κB activation via transgenic expression of constitutively active IKKβ increased the proportion of Tregs among CD4^+^ T cells in the spleen to approximately 30-50% (Long *et al*, 2009). Similarly, mice deficient in the NF-κB negative regulators CYLD or ABIN1 showed enhanced Treg formation, and a similar phenotype was observed in mice with T cell-specific deletion of the NF-κB inhibitor A20 (Fischer *et al*, 2017; Janusova *et al*, 2024; Lee *et al*, 2010; Zhao *et al*, 2011). Interestingly, effector Tregs enriched in *Tank/Azi2*^DKO^ mice express high levels of several TNF receptor superfamily members, including GITR, OX40, 4-1BB, TNFR2, and CD30, all of which can activate the NF-κB pathway (Ward-Kavanagh *et al*, 2016). However, whether enhanced Treg formation in *Tank/Azi2*^DKO^ mice is driven by increased NF-κB signaling in T cells remains to be determined.

TANK and AZI2 recruit TBK1 and IKKε to the TNFR1 signaling complex (Lafont *et al*., 2018). Both kinases were also previously shown to be strongly activated in T cells upon antigenic stimulation (Yu *et al*, 2015), and our data demonstrated that TANK and AZI2 are involved in this pathway. We therefore initially hypothesized that TBK1 and/or IKKε were responsible for suppressing Treg formation. However, this is likely not the case. First, pharmacological inhibition of these kinases did not promote Treg differentiation in vitro. Second, mice deficient in both kinases on the *Ripk1*^KD^ genetic background are viable but do not exhibit increased Treg formation (Eren *et al*., 2024). Altogether, these data indicate that TANK and AZI2 regulate Treg formation independently of their roles in TBK1 and IKKε activation. While ablation of AZI2 alone does not enhance Treg formation, *Tank*^KO^ mice showed a significantly increased proportion of Tregs, which was further elevated in *Tank/Azi2*^DKO^ mice. Since *Tank*^KO^ mice do not develop an overt phenotype, targeting TANK in Tregs might be a strategy to safely promote Treg formation.

Ablation of Tregs leads to severe autoimmunity driven by uncontrolled activation of autoreactive T cells (Lin *et al*, 2005). Therefore, it is not possible to directly assess the role of Tregs in the disease progression of *Tank/Azi2*^DKO^ mice. To overcome this problem, we used mice lacking all T cells, including Tregs. Surprisingly, T cell deficiency exacerbated the phenotype of *Tank/Azi2*^DKO^ mice, indicating that T cells suppress TNFR1- and RIPK1-activity-dependent disease development in this model. Because conventional T cells are unlikely to protect against autoinflammation, and because the majority of T cells in *Tank/Azi2*^DKO^ mice are Tregs, the most plausible explanation is that the exacerbated disease in *Tank/Azi2/Cd3e*^TKO^ mice results from Treg deficiency. Furthermore, we showed that the absence of T cells increases sensitivity to TNF-induced lethality in mice, which is mediated by RIPK1-dependent cell death in nonhematopoietic cells (Zelic *et al*, 2018). These data might likewise be explained by the role of Tregs in suppressing TNF-driven damage.

Although Tregs have not previously been shown to regulate TNFR1-induced cell death, such a function would be consistent with their broader role in maintaining tissue homeostasis. While Tregs are best known for suppressing T cell activity, they can also profoundly influence various processes in the body, including inhibition of proinflammatory cytokine production by innate immune cells, modulation of B cell responses, orchestration of tissue repair and regeneration, and suppression of pain perception (Dikiy & Rudensky, 2023; Midavaine *et al*, 2025; Sage & Sharpe, 2020). Since excessive induction of cell death is highly proinflammatory and leads to immune activation and tissue damage, it seems likely that Tregs have evolved mechanisms to directly limit this pathway.

The hypothesis that Tregs suppress TNF-induced cell death could help explain puzzling findings in SHARPIN-deficient mice, a widely used model of TNF-driven inflammation. SHARPIN is a non-catalytic component of the linear ubiquitin assembly complex (LUBAC), which promotes NF-kB activation and restrains TNF-induced cell death. *Sharpin*^KO^ mice develop chronic lymphoproliferative dermatitis that is rescued by TNF or TNFR1 ablation or by inhibition of RIPK1-dependent cell death pathways (Anderton *et al*, 2022; Berger *et al*, 2014; Gerlach *et al*, 2011; Kumari *et al*, 2014; Peltzer *et al*, 2018; Rickard *et al*, 2014). These mice also exhibit profound defects in Treg development and function (Park *et al*, 2016; Redecke *et al*, 2016; Teh *et al*, 2016). Nevertheless, dermatitis is not driven by the adaptive immune system. Ablation of T and B cells due to RAG1 deletion does not ameliorate skin disease, and transfer of *Sharpin*^KO^ bone marrow into WT recipients fails to induce pathology (HogenEsch *et al*, 1993; Potter *et al*, 2014; Rickard *et al*., 2014). However, *Sharpin*^KO^ mice, which are engineered to express Sharpin only in T cells, have a substantially delayed onset of dermatitis and inflammation, and the effect is even more pronounced when Sharpin is expressed exclusively in Tregs (Sasaki *et al*, 2019). These experiments indicate that restoring functional Tregs in *Sharpin*^KO^ mice can suppress TNF-driven inflammation that is not mediated by conventional T cells.

Altogether, our work shows that T cells in *Tank/Azi2*^DKO^ mice exhibit a markedly increased proportion of Tregs among CD4^+^ T cells, and that T cell ablation exacerbates disease progression in this model. We propose that Tregs restrain inflammation by limiting TNF-induced, RIPK1-dependent cell death. Delineating how T cells and Tregs modulate TNF-induced cell death may open new therapeutic avenues for inflammatory diseases.

## Acknowledgment

This project was supported by the Czech Science Foundation grant (21-25251S) and the Swiss National Science Foundation (320030-227564). OS was supported by the Czech National Health Council (NW26-05-00027). MP and TS were doctoral students supported by the First Faculty of Medicine, Charles University in Prague (SVV 260763). VN and AMM were doctoral students supported by the Faculty of Science, Charles University in Prague. The authors have no conflicting financial interests.

## Author Contributions

MP planned, performed, and analyzed most of the experiments. DP performed the scRNA-seq experiment. DK, AS, TS, AD, and AMM contributed to the analysis of mouse models. VN, JM, and OS contributed to the analysis of scRNA-seq data. OS provided animal models. PD supervised the study and wrote the manuscript. All authors commented on the manuscript draft.

## Methods

### Antibodies

Fluorochrome-conjugated antibodies for flow cytometry against mouse CD3 FITC (#100203), CD4 BV650 (#100545), CD4 AF700 (#100536), CD8α BV421 (#100738), CD25 APC (#102011), CD44 PE (#103007), CD62L BV510 (#104441), CD49d AF647 (#103613), CD357 (GITR) APC (#126311), TNFR2 PE (#113405), CD69 FITC (#104505), HELIOS Pacific Blue (#137210), CD73 PerCP/Cyanine5.5 (#127213), CD278 (ICOS) PE (#107705), CD134 (OX-40) PE (#119409), CD137 (4-1BB) APC (#106109), KLRG1 PerCP/Cyanine5.5 (#138417), CD19 FITC (#152404), CD19 PE (#152407), CD11b BV421 (#101235), TNFR1 APC (#400911), and Ly5.2 AF700 (#109822) were obtained from BioLegend. Anti-mouse Foxp3 PE-Cy7 (#25-5773-80) was purchased from Thermo Fisher Scientific. Anti-Ly5.1 FITC (#553775) was obtained from BD Pharmigen.

Unconjugated antibodies against phospho-TBK1 (S172) (#5483S) and phospho-IKKε (S172) (#8766S) were obtained from Cell Signaling Technology and detected using donkey anti-rabbit IgG AF488 (711-545-152) secondary antibody from Jackson Immunoresearch.

### Mice

Mice were housed in a specific pathogen-free facility with a 12 h/12 h light/dark cycle. Temperature and relative humidity were maintained at 22 ± 1 °C and 55 ± 5 %, respectively, and the mice were provided with a standard rodent breeding diet and water ad libitum. Both males and females were used for experiments. When possible, littermates were equally divided among the experimental groups. All animal protocols were approved by the Resort Professional Commission for Approval of Projects of Experiments on Animals of the Czech Academy of Sciences, Czech Republic. Experimental mice were euthanized by cervical dislocation, isoflurane anesthesia followed by cardiac puncture, or isoflurane inhalation at overdose.

The murine strains harboring *Tank*^KO^, *Azi2*^KO^, *Tnfr1*^KO^, and *Ripk1*^KD^ alleles were previously generated at the Czech Center for Phenogenomics (Ujevic *et al*, 2024). Ly5.1 (RRID: IMSR_JAX:002014), *Cd3e*^KO^ (RRID:IMSR_JAX:004177) (DeJarnette *et al*, 1998), and Rag2^KO^ (RRID:MGI:2174910) (Shinkai *et al*, 1992) strains were described previously. All the mice had a C57Bl/6J background.

### Flow cytometry

Spleens and peripheral lymph nodes were collected from 8-12-week-old mice, and spleens were weighed. Thymi were collected from 23-26-day-old mice. The organs were dissociated into single-cell suspensions using nylon mesh. To remove red blood cells (RBC), splenocytes were incubated for 3 min at room temperature in RBC lysis buffer (eBioscience). Cells were resuspended in FACS buffer (2% fetal bovine serum (FBS), 0.1% sodium azide in PBS) and stained on ice with the LIVE/DEAD near-IR dye (Life Technologies), followed by fixation with eBioscience™ Foxp3 / Transcription Factor Staining Buffer Set (ThermoFisher, cat# 00-55233-00). Fixed samples were stained with the following antibody mixtures. To analyze T cell populations in spleen and lymph nodes, cells were stained with anti-mouse CD3 FITC, CD4 AF700, CD8α BV421, CD25 APC, CD49d AF647, CD44 PE, CD62L BV510, and Foxp3 PE-Cy7 antibodies. To analyze markers of splenic Tregs, splenocytes were stained with antibodies against CD3 FITC, CD4 AF700, Foxp3 PE-Cy7, and GITR APC, 4-1B APC, TNFR2 PE, ICOS PE, or OX40 PE. For the analysis of TNFR1 expression, the cells were stained with anti-mouse CD3 FITC, CD19 PE, CD11b BV421, and TNFR1 APC antibodies. For bone marrow transfer analysis, cells were stained with antibodies against Ly5.2 AF700, Ly5.1 FITC, CD8α BV421 (#100738), CD4 BV650, Foxp3 PE-Cy7, and GITR APC. Samples were measured on the five-laser Cytek Aurora flow cytometer, and data were analyzed using FlowJo software v10.10.0 (BD Biosciences). The gating strategy is shown in Supplementary Figures 1 and 2.

### scRNA sequencing

Spleens from 6-7-week-old mice were collected, and erythrocytes were lysed with RBC lysis buffer for 4 min. Cells were then washed in PBS, resuspended in PBS containing 2% FBS, and stained on ice with the LIVE/DEAD near-IR dye, anti-CD4 PE (#130310) from Biolegend, anti-CD3 FITC (#553062) from BD, and one of the following TotalSeq-C anti-mouse hashtag (MH) antibodies containing anti-CD45 (clone 30-F11), anti-H-2 (clone M1/42) and conjugated to oligonucleotide barcode MH1 (# 155861), MH4 (#155867), MH7 (#155873), MH8 (#155875), MH9 (#155877), or MH10 (#155879) from Biolegend.

Viable CD3^+^ CD4^+^ T cells were sorted using the Influx sorter (BD). Individual samples were then pooled. Cell viability and concentration after sorting were measured using the TC20 Automated Cell Counter (Bio-Rad). The viability of the cells pre-loading was more than 90%. Cells were loaded onto a 10X Chromium machine (10X Genomics) aiming at the yield of 15,000 cells and processed using Chromium Next GEM Single Cell 5′ Reagent Kits v2 (Dual Index) with Feature Barcode technology for CRISPR Screening and Cell Surface Protein (10x Genomics; #PN-1000263, #PN-1000286, #PN-1000541, #PN-1000190, #PN-1000215, #PN-1000250) and Chromium Single Cell Mouse TCR Amplification Kit (#PN-1000254), following the manufacturer’s protocol (#CG000511 Rev C).

### Analysis of scRNA-seq data

The murine reference genome GRCm39 (version 109) was obtained from Ensembl (Harrison *et al*, 2024) and prepared using 10X Cell Ranger 7.1.0 (mkref tool) (Zheng *et al*, 2017). Gene expression and feature barcoding matrices were generated by aligning sequences to the reference using 10X Cell Ranger 7.1.0 (count tool) in paired-end (PE) mode with default parameters. The V(D)J reads were mapped to the murine reference obtained from IMGT (Lefranc, 2011) using 10X Cell Ranger 7.1.0.count mode.

The data were subsequently analyzed using the Seurat 5.0.2 package (Hao *et al*, 2024). Cells with fewer than 200 detected features were removed, and ribosomal and mitochondrial genes were excluded following quantification. V(D)J genes were also removed. Cells that were not clearly labeled by a single hashtag or were labeled by multiple hashtags were excluded. Additionally, cells containing more than two TRA chains, more than one productive TRB chain, or more than two non-productive TRB chains were removed as V(D)J duplicates.

Quality control metrics were assessed, including the number of unique features (nFeature_RNA), total counts (nCount_RNA), and mitochondrial gene percentage (percent.mt). Cells with fewer than 500 detected genes or greater than 10% mitochondrial content were excluded from downstream analysis to remove low-quality cells and potential doublets. Following quality-control filtering, the data were re-normalized and scaled. Principal component analysis (PCA) was performed on the top 1,000 variable features. Based on elbow plot inspection, the first 20 principal components were selected for downstream analysis. Uniform Manifold Approximation and Projection (UMAP) was performed using the first 15 PCs for visualization. Cell clustering was performed using a shared nearest neighbor (SNN) modularity optimization-based clustering algorithm at multiple resolutions (0.2-1.2). Optimal clustering resolution (0.4) was determined using Clustree analysis to assess cluster stability. Cluster-specific marker genes were identified using the FindAllMarkers function with the default Wilcoxon rank-sum test, filtering for genes with average log2 fold-change > 1 and expressed in at least 25% of cells within the cluster. Clusters were manually annotated based on canonical T cell marker expression patterns. Cell type frequencies were calculated as percentages within each mouse sample.

### Bone marrow transfer

Bone marrow cells were isolated from 4–5-week-old *Tank/Azi2*^DKO^ mice carrying the Ly5.2 allele and from WT mice expressing Ly5.1/5.2. Cells were mixed in a 1:1 ratio, and 2×10^6^ cells in 200 µl PBS were intravenously injected into lethally irradiated (6 Gy) 6-week-old Ly5.1 recipient mice. After 12–14 weeks, spleens and lymph nodes were harvested and analyzed by flow cytometry.

### Analysis of TBK1 and IKK**ε** activation upon stimulation

Spleens were collected from 8-12-week-old mice and dissociated into single-cell suspensions using a nylon mesh. Splenocytes were incubated in ACK lysis buffer for 3 min at room temperature to remove red blood cells. Cells were resuspended in Iscove’s Modified Dulbecco’s Medium (IMDM) (Lonza Bioscience) medium supplemented with 10% FBS (Biosera) and were left unstimulated or activated with PMA (5 ng/ml) (Sigma-Aldrich, #P1585) and ionomycin (0.5 µM) (Sigma-Aldrich, #I3909) for 15 min at 37 °C. Cells were fixed, permeabilized, blocked with TruStain FcX PLUS (Biolegend, #156603), and subsequently stained with rabbit antibodies against pTBK1 and pIKKε, washed, and stained with an antibody mixture containing anti-CD4 AF700, anti-CD8 BV421, anti-Foxp3 Pe-Cy7, anti-CD19 FITC, and anti-rabbit IgG AF488. Samples were analyzed on the Cytek Aurora flow cytometer.

### Induction of Tregs in vitro

Spleens and peripheral lymph nodes were collected from 8-12-week-old mice, organs were dissociated into single-cell suspensions using nylon mesh, and splenocytes were incubated in ACK lysis buffer for 3 min at room temperature to remove red blood cells. Primary T cells were isolated using the MojoSort™ Mouse CD3 T Cell Isolation Kit (Biolegend, #480031) according to the manufacturer’s protocol. Cells were resuspended in advanced RPMI 1640 medium (Sigma) supplemented with 2 mM L-Glutamine, 1 mM sodium Pyruvate, 10mM HEPES, 1x MEM non-essential amino acids (Thermo Fisher Scientific), 10% FBS and1% penicillin-streptomycin (Biosera) and stimulated with recombinant murine IL-2 (PeproTech, #4501) and anti-CD3/CD28 beads (ThermoFisher #11453D) at a beads-to-cells ratio of 1:1. Where indicated, following inhibitors at a concentration 1 µM were added: Compound 1 (Selleckchem, #S8920), Amlexanox (Cayman chemical, #14181), MRT67307 (Tocris Bioscience, #5134), GSK8612 (Sigma-Aldrich, #SML2721), Rapamycin (Sigma-Aldrich, #553211). Cells were incubated for 5 days at 37 °C, 5% CO2, fixed and permeabilized, stained with CD4 AF700 and Foxp3 Pe-Cy7 antibodies, and analyzed on the Cytek Aurora flow cytometer.

### Histological analysis

Tissue samples from the kidneys of 3-4-week-old mice were fixed in 4% formaldehyde solution in PBS overnight and transferred to 70% ethanol. The samples were processed by the Leica ASP6025 Vacuum Tissue Processor according to the program Standard processing overnight, embedded in Paraplast X-tra (Leica Biosystems), using the Tissue Embedding Station Leica EG1150. Five μm-thick sections were prepared using Leica Fully Motorized Rotary Microtome RM2255-FU. The obtained slides were stained by Hematoxylin H (Biognost) and Eosin Y (Carl Roth) using the Leica ST5010-CV5030 Stainer Integrated Workstation.

### Serum cytokine analysis

To collect blood samples, the mice were deeply anesthetized with isoflurane, and blood was collected via cardiac puncture into EDTA-coated tubes. Blood samples were centrifuged (1200 g, 15 minutes, 4 °C) and plasma samples were frozen at −80 °C until further analysis. The concentration of cytokines in the plasma was measured using the ProcartaPlex mouse cytokine & chemokine panel 1A, 36-plex (Thermo Fisher Scientific, #EPX360-26092-901) on the Bio-Plex 200 system (Bio-Rad).

### TNF production

Murine TNF ligand (amino acids 80-235) was fused at the N-terminus to the CD33 leader and 6xHis Tag, which enabled the secretion and purification of the protein. The coding sequence was prepared using the GeneArt Gene Synthesis service (Thermo Fisher Scientific) and inserted into the pcDNA3.1 vector. The plasmid was transfected into HEK293FT cells using polyethylenimine (PEI) transfection. After three days, cell supernatants containing the protein of interest were collected and loaded on a His GraviTrap TALON column (Cytiva), previously equilibrated with a purification buffer (50 mM sodium phosphate, pH 7.4, 300 mM NaCl). The columns were washed with a purification buffer containing 20 mM imidazole and eluted with a purification buffer containing 350 mM imidazole. Subsequently, the sample was loaded on a centrifugal filter (10 kDa molecular weight cut-off, Millipore), washed several times with purification buffer to remove imidazole, and concentrated. The concentration of purified protein was determined using NanoDrop One (Thermo Fisher Scientific). Concentrated ligand was supplemented with 50% glycerol and kept at −80 °C for long-term storage.

### Intravenous TNF administration

Mice were weighed before the experiment and subsequently injected with TNF at a concentration of 1 µg per 1 g of body weight in 150 µl of PBS via the lateral tail vein. Animals were monitored at one or two-hour intervals. Rectal temperature was measured using a lubricated digital thermometer to assess systemic responses. Mice exhibiting a body temperature below 26 °C were euthanized. Where indicated, 30 min prior to TNF injection, mice were first administered with an intraperitoneal injection of 200 µg of the RIPK1 inhibitor Nec-1s per 20 g of body weight, dissolved in 200 µl of PBS or injected with vehicle alone.

## Supplementary Figures

**Supplementary Figure 1.**
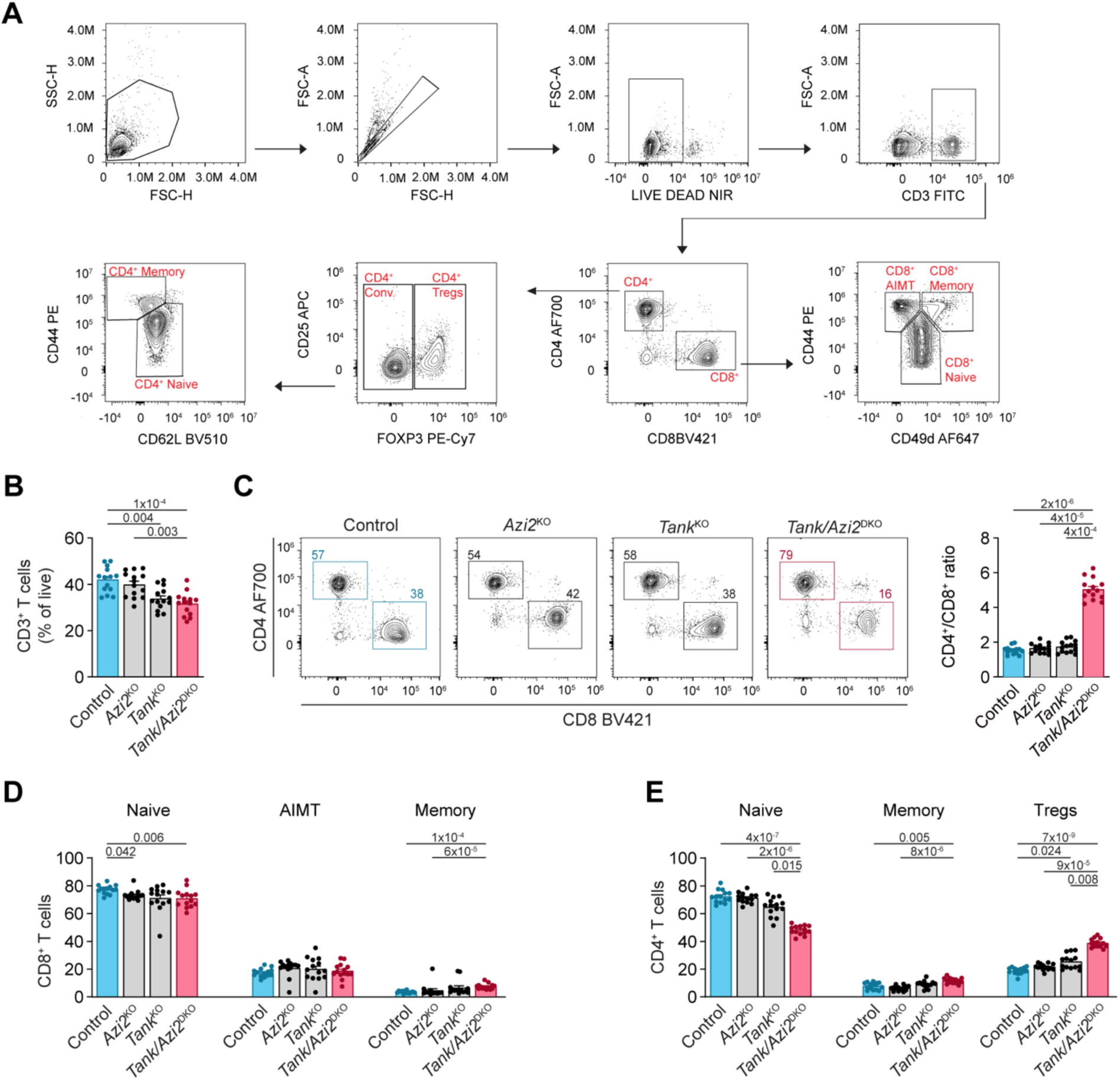
*Tank* and *Azi2* suppress the formation of Tregs. (**A**) Gating strategy used for analysis of T cells isolated from the spleen (Fig. 1) and peripheral lymph nodes (Fig. 1S). (**B-E**) Flow cytometry analysis of T cells isolated from peripheral lymph nodes of 8-12-week-old mice. (B) Proportion of live T cells (gated as CD3^+^). (C) Ratio of CD4^+^ to CD8^+^ T cells. (D) Fraction of CD8^+^ T cells gated as naïve (CD44^−^), memory (CD44^+^, CD49d^+^), and antigen-inexperienced memory T (AIMT) cells (CD44^+^, CD49d^−^). (E) Fraction of CD4^+^ T cells gated as Tregs (FOXP3^+^), naïve (FOXP3^−^, CD44^−^, CD62L^+^), and memory (FOXP3^−^, CD44^+^, CD62L^−^). Data in (B-F) are presented as mean + SEM (n = 14 mice per group), Kruskal–Wallis test with Dunn’s multiple-comparisons post-hoc test.

**Supplementary Figure 2.**
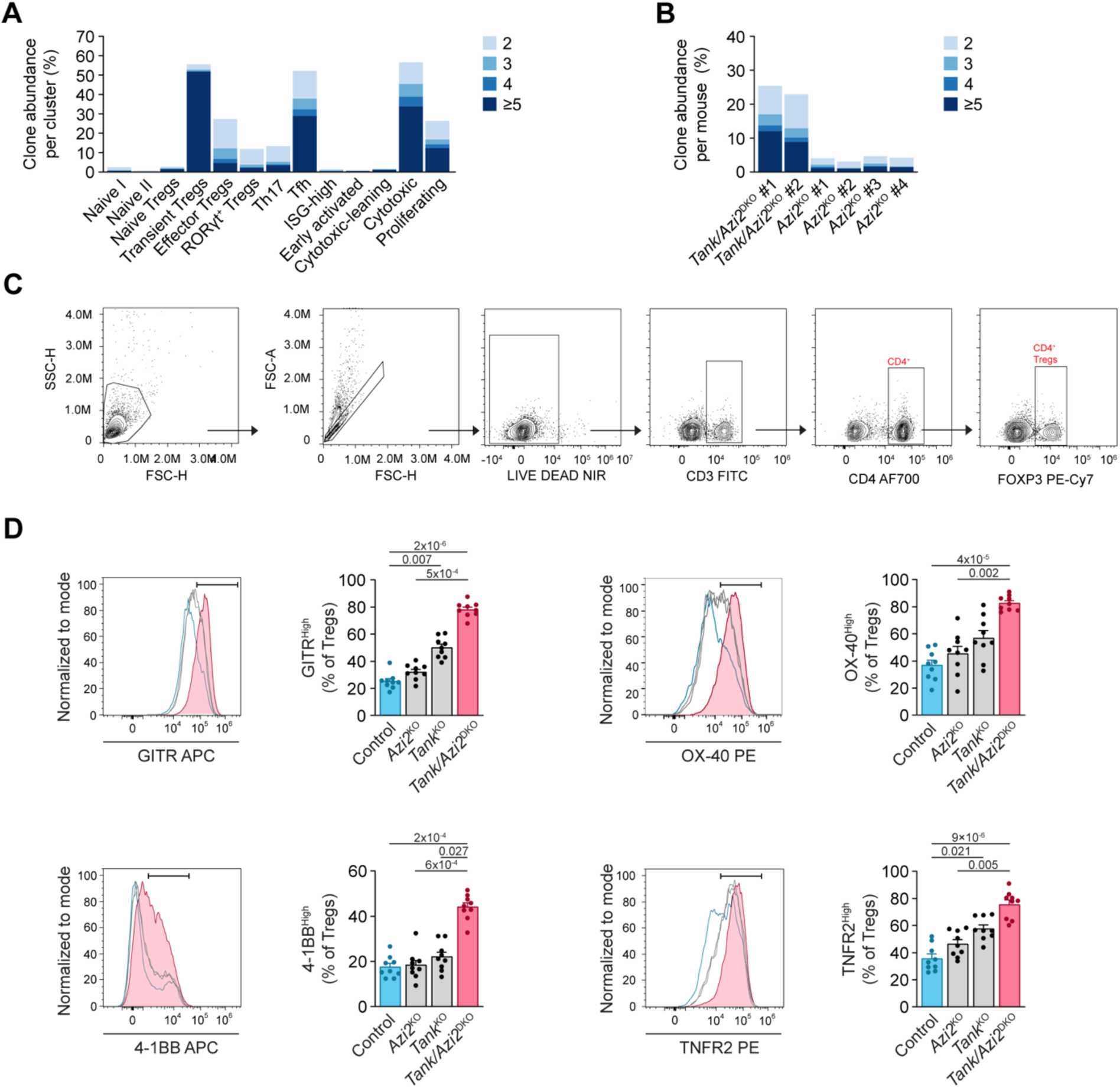
The majority of Tregs in *Tank/Azi2*^DKO^ mice exhibit an effector phenotype. (**A-B**) Clonal abundance of CD4^+^ T cells in each cluster (A) and in individual experimental mice (B), as determined by V(D)J sequencing from scRNA-seq analysis. (**C-D**) Flow cytometry analysis of splenic Tregs from 8 to 12-week-old mice. (C) Gating strategy used to identify Tregs. (D) Expression of the indicated activation markers on Tregs. Bars mark the gate defining cells with high expression of each marker. Data are presented as mean + SEM (n = 9 mice per group). Kruskal–Wallis test with Dunn’s multiple-comparisons post hoc test.

**Supplementary Figure 3.**
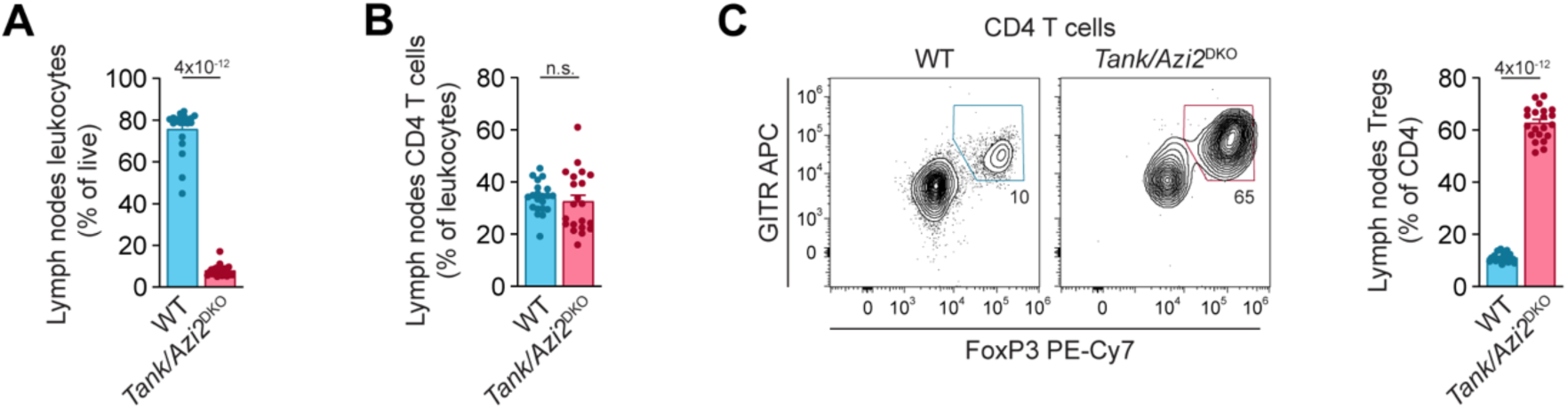
*Tank/Azi2*^DKO^ CD4^+^ T cells have an intrinsic propensity to form Tregs at the periphery. (**A-C**) Flow cytometry analysis of T cells isolated from peripheral lymph nodes of irradiated recipient mice reconstituted with mixed bone marrow chimeras generated from WT and *Azi2/Tank*^DKO^ donors (n = 21 recipient mice in 3 separate experiments). (A) Percentage of lymph node leukocytes derived from the indicated bone marrow donor cells. (B) Percentage of CD4⁺ T cells among lymph node leukocytes. (C) Frequency of Tregs (GITR^high^, FOXP3⁺) within the CD4⁺ T cell population. Data are shown as mean + SEM. Two-tailed Mann-Whitney test. n.s., not significant.

**Supplementary Figure 4.**
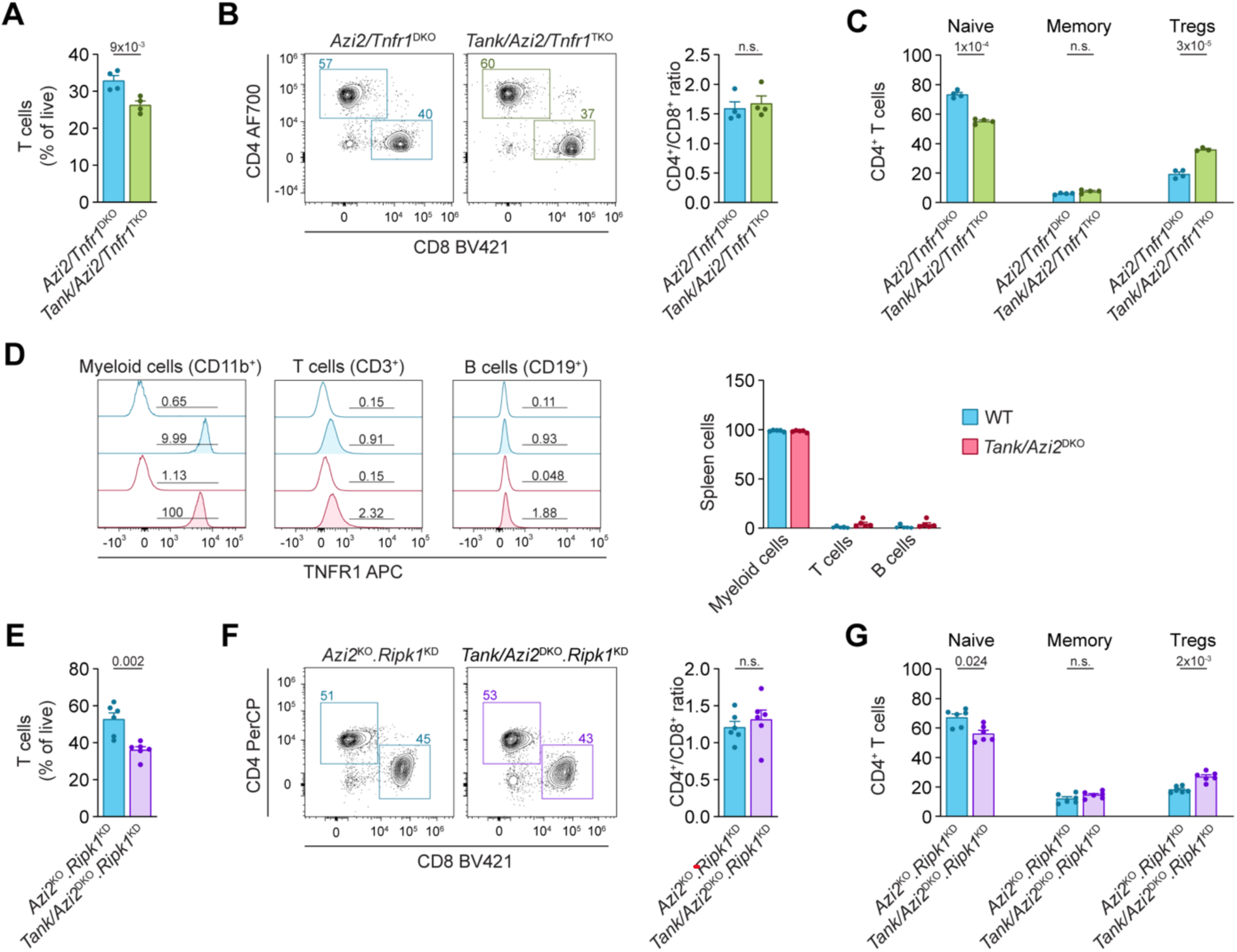
Enhanced Tregs formation in *Tank/Azi2*^DKO^ mice is partially driven by TNFR1-mediated inflammation. (**A-C**) Flow cytometry analysis of T cells isolated from peripheral lymph nodes of 8-12-week-old *Azi2/Tnfr1*^DKO^ and *Tank/Azi2/Tnfr1*^TKO^ mice (n = 4 mice per group). (A) Proportion of live T cells (gated as CD3^+^). (B) Ratio of CD4^+^ to CD8^+^ T cells. (C) Fraction of CD4^+^ T cells gated as Tregs (FOXP3^+^), naïve (FOXP3^−^, CD44^−^, CD62L^+^) and memory cells (FOXP3^−^, CD44^+^, CD62L^−^). (**D**) Flow cytometry analysis of TNFR1 expression on myeloid cells (*CD11b*^+^), T cells (*CD3*^+^), and B cells (*CD19*^+^) isolated from the spleen of 8-12-week-old WT and *Tank/Azi2*^DKO^ mice (n = 6 mice per group). (**E-G**) Flow cytometry analysis of T cells isolated from peripheral lymph nodes of 8-12-week-old *Azi2*^KO^.*Ripk1*^KD^ and *Tank/Azi2*^DKO^.*Ripk1*^KD^ mice (n = 6 mice per group). Cells were analyzed as in (A-C). Data are presented as mean + SEM. Two-tailed Mann-Whitney test.

**Supplementary Figure 5.**
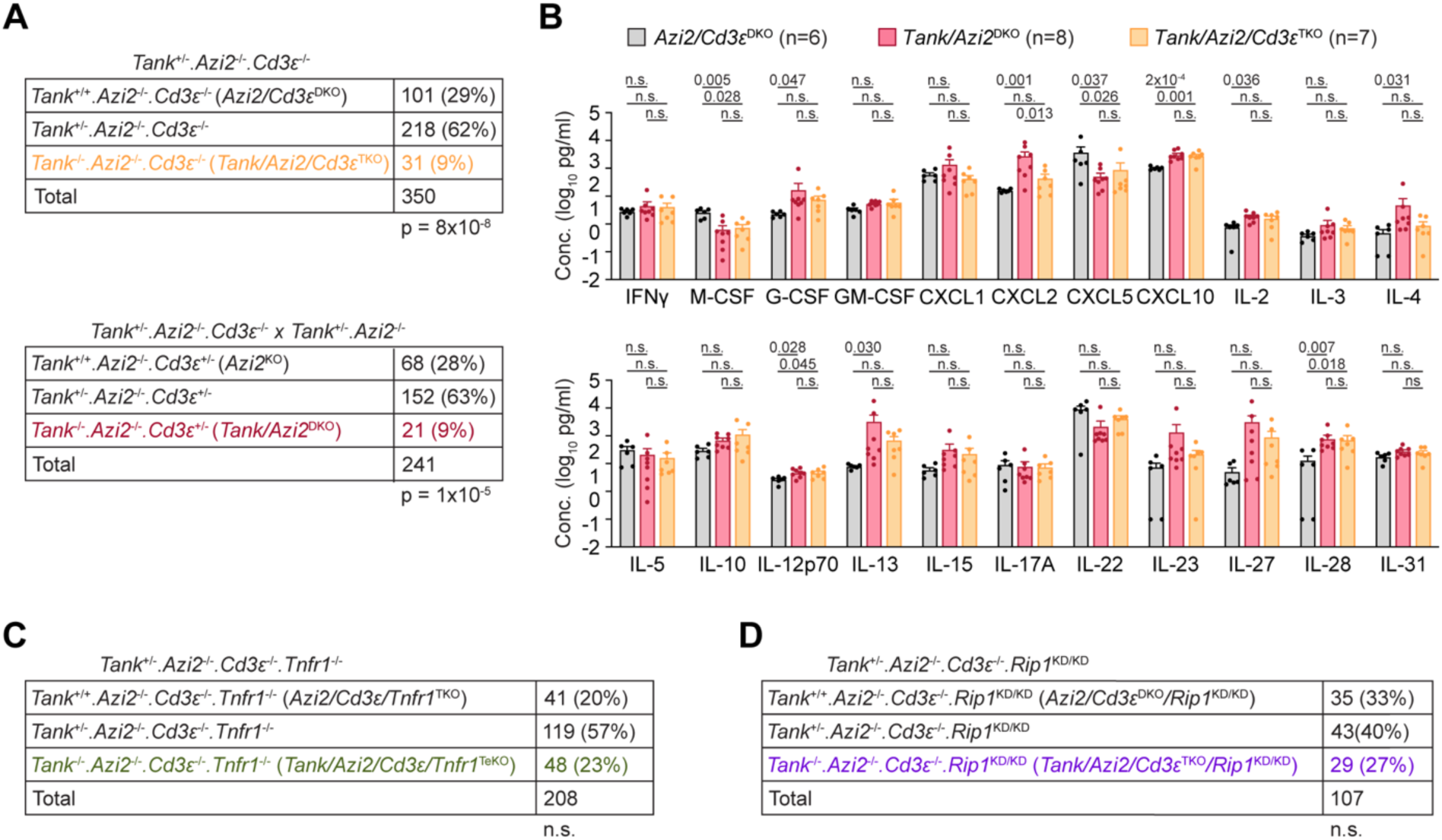
T cell ablation exacerbates TNF-driven autoinflammatory disease in *Tank/Azi2*^DKO^ mice. (**A, C, D**) Genotypes and numbers of pups born to parents of the indicated genotypes, chi-square test. (**B**) Plasma cytokine concentrations in 22-26-day-old mice. Mean + SEM. One-way ANOVA with Tukey’s post-tests. n.s., not significant.

**Supplementary Figure 6.**
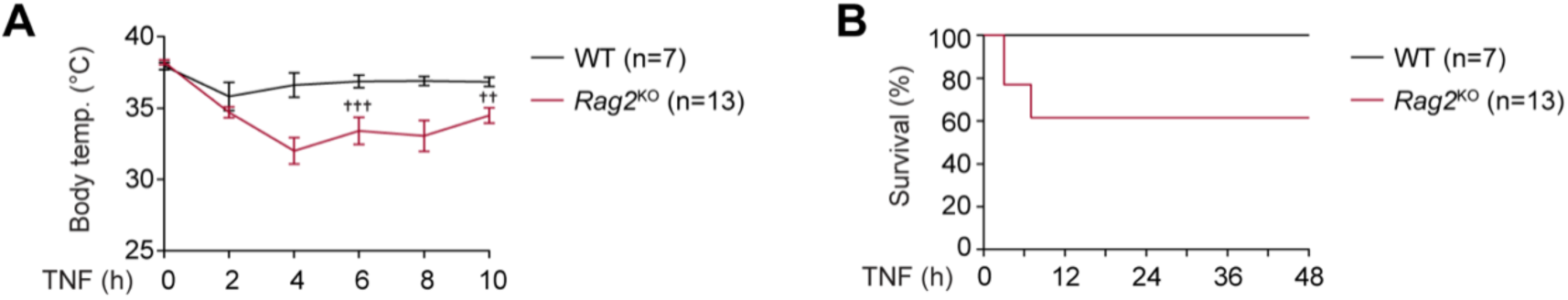
Adaptive immune cells protect against TNF-induced cell death. (**A-B**) WT (n = 7) or *Rag2*^KO^ (n = 13) littermates were injected with TNF (1 µg/g body weight). (A) Body temperature was measured every 2 h for up to 10 h. (B) Proportion of mice reaching humane endpoint. The mice that reached humane endpoint are indicated by a cross.

